# SARS-CoV-2 Infection of Salivary Glands Compromises Oral Antifungal Innate Immunity and Predisposes to Oral Candidiasis

**DOI:** 10.1101/2024.05.13.593942

**Authors:** Areej A. Alfaifi, Tristan W. Wang, Paola Perez, Ahmed S. Sultan, Timothy F. Meiller, Peter Rock, David E. Kleiner, Daniel S. Chertow, Stephen M. Hewitt, Billel Gasmi, Sydney Stein, Sabrina Ramelli, Daniel Martin, Blake M. Warner, Mary Ann Jabra-Rizk

## Abstract

Saliva contains antimicrobial peptides considered integral components of host innate immunity, and crucial for protection against colonizing microbial species. Most notable is histatin-5 which is exclusively produced in salivary glands with uniquely potent antifungal activity against the opportunistic pathogen *Candida albicans*. Recently, SARS-CoV-2 was shown to replicate in salivary gland acinar cells eliciting local immune cell activation. In this study, we performed mechanistic and clinical studies to investigate the implications of SARS-CoV-2 infection on salivary histatin-5 production and *Candida* colonization. Bulk RNA-sequencing of parotid salivary glands from COVID-19 autopsies demonstrated statistically significant decreased expression of histatin genes. *In situ* hybridization, coupled with immunofluorescence for co-localization of SARS-CoV-2 spike and histatin in salivary gland cells, showed that histatin was absent or minimally present in acinar cells with replicating viruses. To investigate the clinical implications of these findings, salivary histatin-5 levels and oral *Candida* burden in saliva samples from three independent cohorts of mild and severe COVID-19 patients and matched healthy controls were evaluated. Results revealed significantly reduced histatin-5 in SARS-CoV-2 infected subjects, concomitant with enhanced prevalence of *C. albicans*. Analysis of prospectively recovered samples indicated that the decrease in histatin-5 is likely reversible in mild-moderate disease as concentrations tended to increase during the post-acute phase. Importantly, salivary cytokine profiling demonstrated correlations between activation of the Th17 inflammatory pathway, changes in histatin-5 concentrations, and subsequent clearance of *C. albicans* in a heavily colonized subject. The importance of salivary histatin-5 in controlling the proliferation of *C. albicans* was demonstrated using an *ex vivo* assay where *C. albicans* was able to proliferate in COVID-19 saliva with low histatin-5, but not with high histatin-5. Taken together, the findings from this study provide direct evidence implicating SARS-CoV-2 infection of salivary glands with compromised oral innate immunity, and potential predisposition to oral candidiasis.

**AUTHOR SUMMARY:** Saliva contains antimicrobial peptides part of host innate immunity crucial for protection against colonizing microbial species. Most notable is the antifungal peptide histatin-5 produced in salivary glands cells. SARS-CoV-2 was shown to replicate in salivary gland cells causing tissue inflammation. In this study, we showed decreased expression of histatin genes in salivary glands from COVID-19 autopsies, and co-localization studies of SARS-CoV-2 spike and histatin revealed absence or minimal presence of histatin in acinar cells with replicating virus. To investigate the clinical implications of these findings, we conducted studies using saliva samples from subjects with mild to severe COVID-19, matched with healthy controls. Results revealed significantly reduced histatin-5 in SARS-CoV-2 infected subjects with enhanced prevalence of *C. albicans*. Prospective analysis indicated the decrease in histatin-5 is reversible in mild-moderate disease, and salivary cytokine profiling demonstrated activation of the Th17 inflammatory pathway. The importance of salivary histatin-5 in controlling the proliferation of *C. albicans* was demonstrated using an *ex vivo* assay where *C. albicans* was able to proliferate in saliva with low histatin-5, but not with high histatin-5. Collectively, the findings provide direct evidence implicating SARS-CoV-2 infection of salivary glands with compromised oral innate immunity and predisposition to oral candidiasis.

## INTRODUCTION

The oral cavity remains an underappreciated site for SARS-CoV-2 infection despite the evident myriad of oral conditions observed in COVID-19 patients, and the presence of the virus in saliva (1–6). The homeostasis of the oral cavity is maintained by saliva, an extracellular fluid produced by salivary glands possessing a wealth of protective properties (7). Specifically, saliva is enriched with antimicrobial peptides considered to be part of the host Th17-type adaptive immune response that play a vital role in innate immune defenses against microbial species (7–10). Most notable are histatins, a family of peptides exclusively produced in the acinar cells of the salivary glands and secreted into saliva (11, 12). There are two genes for histatins, *HTN1* and *HTN3* which encode histatins-1 and 3, respectively (13). Histatin-5, a proteolytic product of histatin-3, is the most abundant and unique as it exhibits potent antifungal activity against the fungal pathogen *Candida albicans* (*C. albicans*) (12, 14–18). Although *C. albicans* is a commensal colonizer of the oral cavity, any changes in the host microenvironment favoring its proliferation allows this opportunistic species to transition into a pathogen causing oral candidiasis (thrush) (19, 20). As a commensal colonizer of the oral mucosa, the host immune response to *Candida* is oriented toward a more tolerogenic state and, therefore, local innate immune defenses and specifically histatin-5 play a central role in maintaining *Candida* in its commensal state preventing development of oral candidiasis (8, 21, 22).

Recently, the salivary glands were shown to be a potential target for SARS-CoV-2 infection as several studies demonstrated the concomitant expression of ACE2/transmembrane serine proteases 2 (TMPRSS2) in salivary glands epithelial cells (23–27). In fact, virus entry into salivary glands cells was found to be higher, as compared with entry into lung cells (25). Most notable are findings from a landmark study by Huang and Perez *et al.* (2021) (28) which comprehensively demonstrated that salivary gland acinar and ductal cells are robust sites for SARS-CoV-2 infection and replication, eliciting local immune cell activation. Additionally, architectural distortion, atrophy, fibrosis, and ductal rupture were also revealed, establishing that the minor and major salivary glands are susceptible sites for infection, replication and local immune cell activation. Moreover, the authors demonstrated that saliva from acutely infected patients could infect and replicate in Vero cells, *ex vivo*, indicating that the source of virus in saliva is likely derived from infected cells in the salivary glands (28). In fact, clinical observations are in support of SARS-CoV-2 mediated damage to the salivary glands as COVID-19 patients frequently present with gustatory dysfunction and xerostomia, and cases of inflamed salivary glands have been reported in this population (26, 29).

Unlike other antimicrobial peptides, histatins are exclusively expressed and secreted by acinar cells of salivary glands; saliva containing histatins is secreted from the acini into the lumen which transits the ductal network into the oral cavity where they can exert their antimicrobial activities. Patients infected with SARS-CoV-2 have the propensity to develop superimposed infection including oral candidiasis. While SARS-CoV-2 infects and replicates in the acinar and ductal cells of the salivary glands, it is not known if the risk of oral candidiasis is due to affects on the acinar expression of anti-candidal proteins, or due to viral and immune-mediated destruction of the glands. To that end, we performed mechanistic and clinical studies to investigate the implications of SARS-CoV-2 infection of salivary gland tissue on oral innate immune defenses, salivary histatin-5 production, and predisposition to opportunistic infections. Using *in situ* hybridization and immunofluorescence, we performed co-localization studies to assess histatin production in SARS-CoV-2 infected salivary gland acinar cells. To provide mechanistic insights, the expression of histatin genes was comparatively evaluated in infected and uninfected salivary gland tissue. The clinical implications of findings from mechanistic studies were demonstrated using various cohorts of SARS-CoV-2 infected subjects to evaluate salivary histatin-5 levels and oral *Candida* colonization. Collectively, the novel findings from this study establish the oral cavity as a robust site for SARS-CoV-2 infection warranting reassessment of the risks for oral opportunistic infections in COVID-19 patients.

## RESULTS

### SARS-CoV-2 infection of the parotid glands affects the expression of histatins

To determine the effect of SARS-CoV-2 infection on the expression of anti-candidal proteins in the salivary glands, RNA sequencing (RNAseq) and confirmatory hybrid *in situ* hybridization and immunofluorescence microscopy, was used. First, RNAseq of SARS-CoV-2 infected parotid glands showed significantly reduced expression of histatin genes (*HTN1,* >100-fold lower, p=0.0022; *HTN3,* >100-fold lower, p=0.0038; Fig. 1a). Next, we confirmed the loss of histatin expression and the co-localization of SARS-CoV-2 and histatin protein expression in acinar cells. Hybrid *in situ* hybridization and immunofluorescence microscopy co-localization studies of SARS-CoV-2 and histatins, respectively, demonstrated significant (p<0.001) reduction of histatin protein expression in virus infected acinar cells in parotid gland tissues obtained from deceased COVID-19 patients. As previously reported, the detection of SARS-CoV-2 is non-uniform across the acini of the glands (28). Comparing infected vs non-infected acinar cells, histatin protein expression intensity was inversely proportional to viral count (Fig.1b-f) indicating that direct viral infection of the glands suppresses histatin mRNA and protein expression.

**Figure 1.**
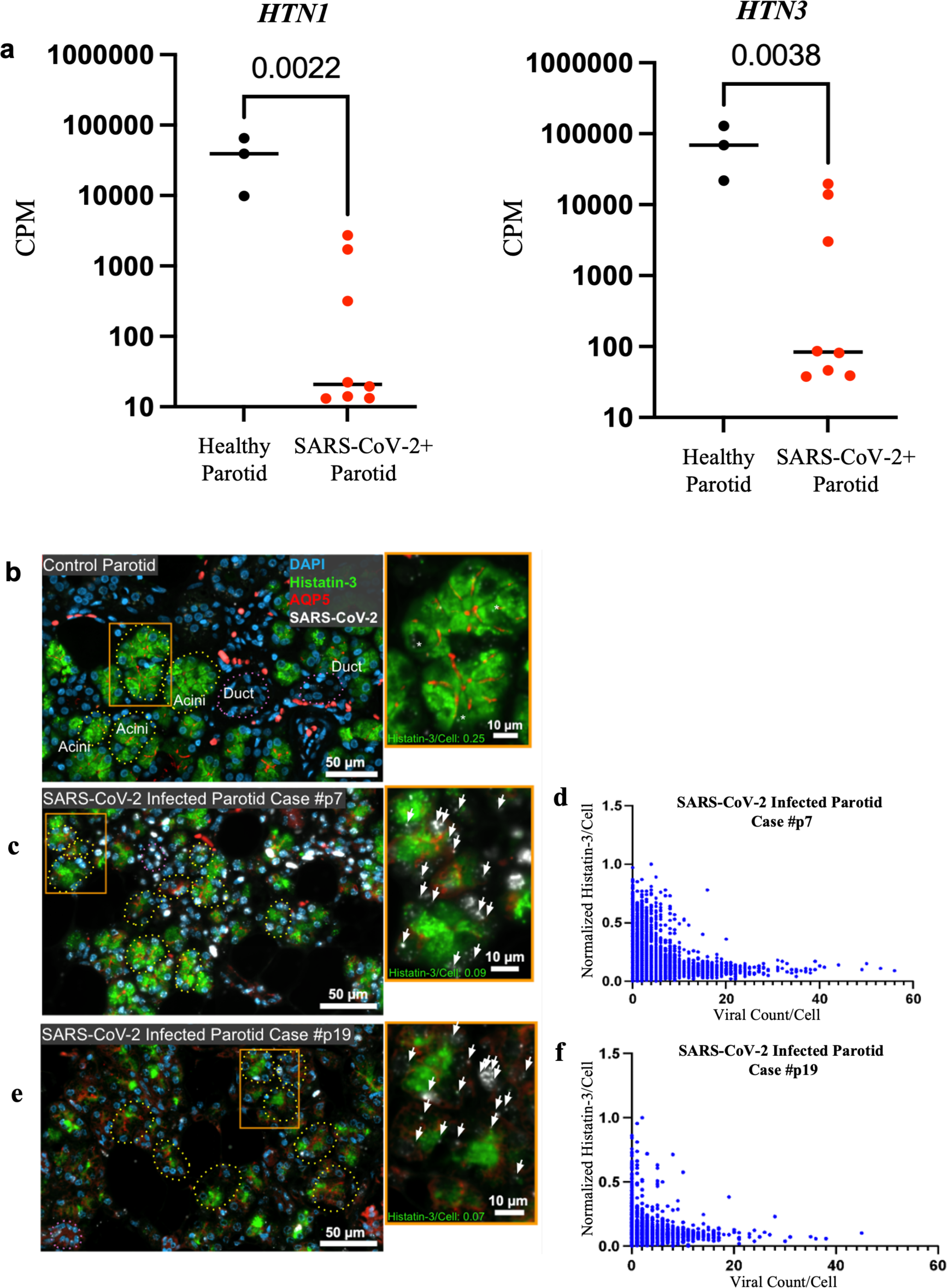
Impact of SARS-CoV-2 salivary gland infection on histatin production, gene expression and salivary levels. **(a)** Parotid expression of histatin genes (*HTN1* and *HTN3*) from deceased COVID-19 subjects (n=8) and healthy control subjects (n=3) using bulk RNA sequencing. **(b-f)** Co-localization studies using *in situ* hybridization for SARS-CoV-2 (*white dots* and *arrows*) and immunofluorescence for histatin-3 (*green*) with quantitative correlation between the intensity of histatin and SARS-CoV-2 counts within infected acinar cells of parotid tissue from **(b)** healthy subject (*white asterisk,* non-specific signal) and **(c, e)** deceased COVID-19 patients. Acini (dotted circles) identified based on expression of AQP5 (*red,* apical membrane); *inset* (*orange box*) shows the presence of the viral genome in acinar cells of the parotid glands. **(d, f)** Per acinar cell expression of histatin and the per cell viral count are inversely proportional as shown in two representative COVID-19 cases (P7, P19).

### SARS-infected patients secrete significantly less histatins potentiating oral candidiasis

Clinical studies were conducted using two COVID-19 patient cohorts (hospitalized [UMD] and outpatient [NIDCR]) with ranging disease severity, and matched healthy controls. In the hospitalized cohort, a histatin-5 specific immunoassay to measure salivary histatin-5 concentrations was used and significant (p=0.0303) differences in values between the COVID-19 patients and matched healthy controls were seen. Average concentrations for the control group was 21.3 µg/ml (11.1 µg/ml-34.5 µg/ml), and 18.4 µg/ml (1.4 µg/ml-53.5 µg/ml) for the COVID-19 group. Strikingly, 14 of the COVID-19 patients had concentrations below 10 µg/ml (Fig. 2a), and although unclear why, 3 subjects had concentrations higher than any of the controls. The difference in concentrations between the two groups was also seen in age- and race-matched comparison between each COVID-19 patient and their matched control subject (Fig. 2b). *Candida* colonization was assessed by culturing samples; where no *Candida* was recovered from any of the healthy controls, 45% (9/20) of COVID-19 patients sampled were positive for *C. albicans*, some heavily colonized (Fig. 2a).

**Figure 2.**
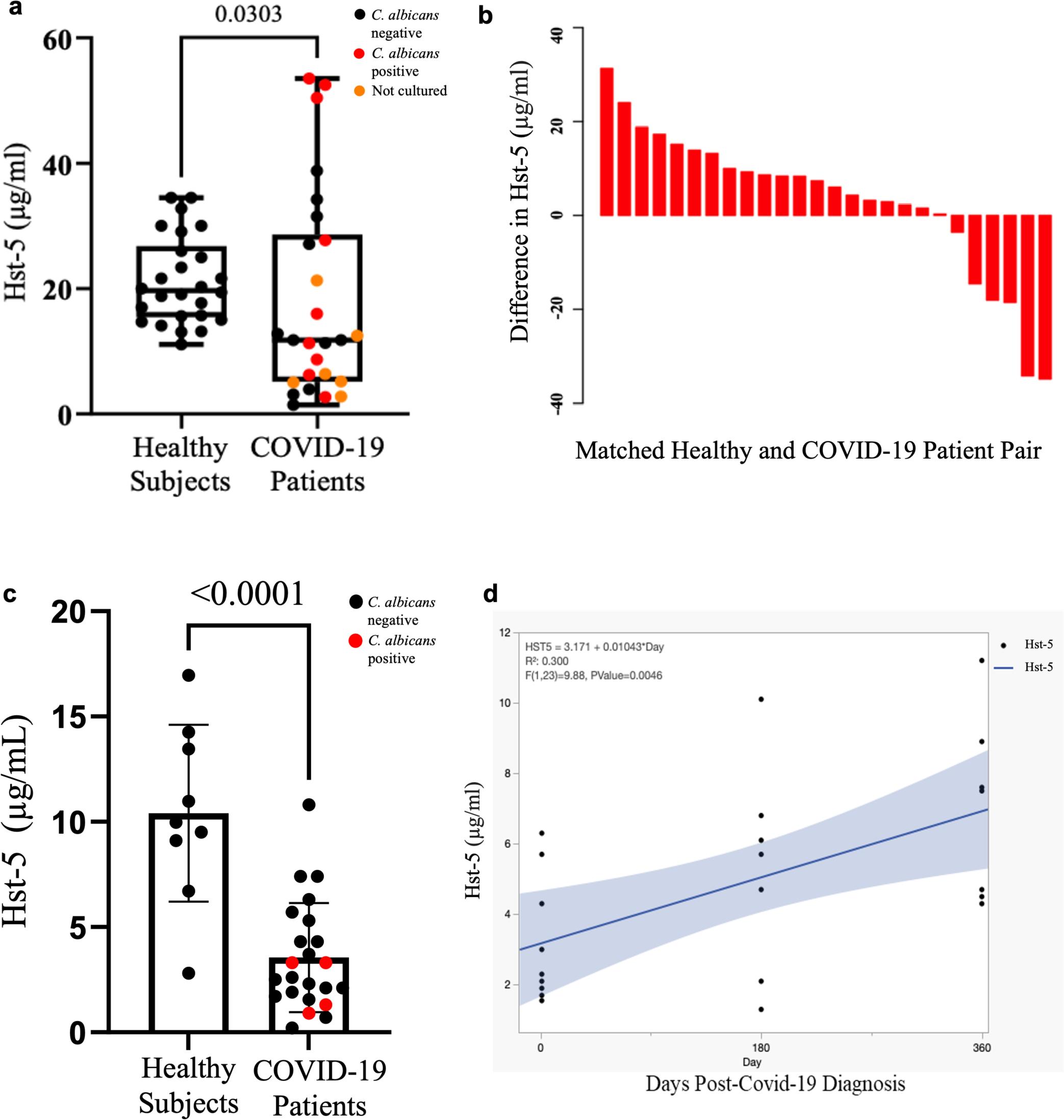
Salivary histatin-5 (Hst-5) levels and fungal colonization in prospectively sampled hospitalized and outpatient COVID-19 cohorts. **(a)** Boxplot with Hst-5 concentrations and *Candida* recovery from 26 saliva samples from hospitalized COVID-19 patients and matched healthy controls. **(b)** Waterfall plot depicting 26 pairs of COVID-19 and healthy subjects matched for race and age. Bar height represents differences among pairs in Hst-5 levels (µg/ml) between healthy controls and COVID-19 patients. **(c)** Bar plot depicts Hst-5 concentrations and *Candida* recovery of 9 healthy subjects and 23 COVID-19 outpatients’ saliva. Red marked dots indicate cultured saliva contained candidal outgrowth **(d)** Linear regression analysis of serially sampled COVID-19 outpatients’ saliva (n=8) shows time-dependent restoration of Hst-5 concentration from the post-acute phase to the chronic phase, based on longitudinally collected samples from acute phase (3-15 days) to chronic phase (6-12 months). Blue line, linear fit; light blue shading, confidence interval of the linear fit.

To corroborate these findings, a separate outpatient cohort collected by the National Institute of Dental and Craniofacial Research (NIDCR) (28) of serially sampled acutely infected patients was analyzed for histatin-5 expression. Compared to healthy control samples collected prior to the pandemic (n=9), the acute phase COVID-19 whole unstimulated saliva (n=23) had significantly lower histatin-5 concentrations (3.63 vs 10.41 µg/ml, *p*<0.0001; Fig. 2c). *Candida* colonization was assessed as above; no *Candida* was recovered from the healthy controls, yet 17% (4/23) of COVID-19 patients sampled were positive for *C. albicans.* Despite high variability in the expression of histatin-5 across the cohort, prospective analysis of available serially-collected samples (n=8; 3-15 days from symptom onset, 6-months, and 1-year from initial infection) from these COVID-19 subjects indicated a slight, but statistically significant trend in the restoration histatin-5 concentration from the post-acute to the chronic phase (R^2^=0.30, *p*=0.0046; Fig. 2d). However, some patients exhibited persistence of low salivary histatin concentration for up to 1 year after recovery in the chronic phase. No statistical differences (p values > 2.0) in histatin-5 concentrations were seen between subjects based on disease severity or other variables (gender, race, age).

### Significant changes in histatin-5 concentrations over the course of COVID-19 disease in prospectively sampled subjects

A total of 5 subjects with mild-moderate disease were prospectively sampled and salivary histatin-5 was measured to monitor changes in concentrations. Two of the subjects were longitudinally sampled up to 39 and 80 days, respectively (Fig. 3). For these subjects, saliva and histatin-5 concentrations prior to COVID-19 infection were available. The baseline histatin-5 concentration measured prior to COVID-19 disease for *Subject #1* was 15.7 µg/ml. However, the concentration in the first sample recovered during the acute phase of the disease was 5.6 µg/ml (64.33% drop) and 8.8 µg/ml for the second sample recovered 2 days later. The concentration gradually began to increase during the post-acute phase with a spike noted on Day 15 (25.3 µg/ml) returning to baseline level in the last sample analyzed (Fig. 3a). For *Subject #2*, the histatin-5 concentration measured prior to COVID-19 disease was 32.8 µg/ml; however, the concentration in the first sample recovered during the post-acute phase of the disease was 15.65 µg/ml (52.29% drop) which gradually increased over subsequent days with a spike noted on Day 45 (43.6 µg/ml) before returning to pre-COVID-19 levels on last day sampled (Fig. 3c). Significantly, *C. albicans* was recovered from the initial 6 samples recovered from *Subject #2*, but not from the last 3 samples that followed the restoration of histatin-5 to pre-COVID-19 level (Fig. 3c). Three additional subjects (Subjects 3, 4, and 5) were also prospectively sampled (Fig. 4), however, for this group, pre-COVID-19 saliva was not available. For these subjects, the histatin-5 concentrations in the first samples recovered during the acute phase of the disease were 2.4, 4.1 and 5.6 µg/ml, respectively which increased during the post-acute phase to 10.9, 13.2, and 12.5 µg/ml, respectively in the last samples tested (Fig. 4). No *Candida* was recovered from any of the samples from these 3 subjects.

**Figure 3.**
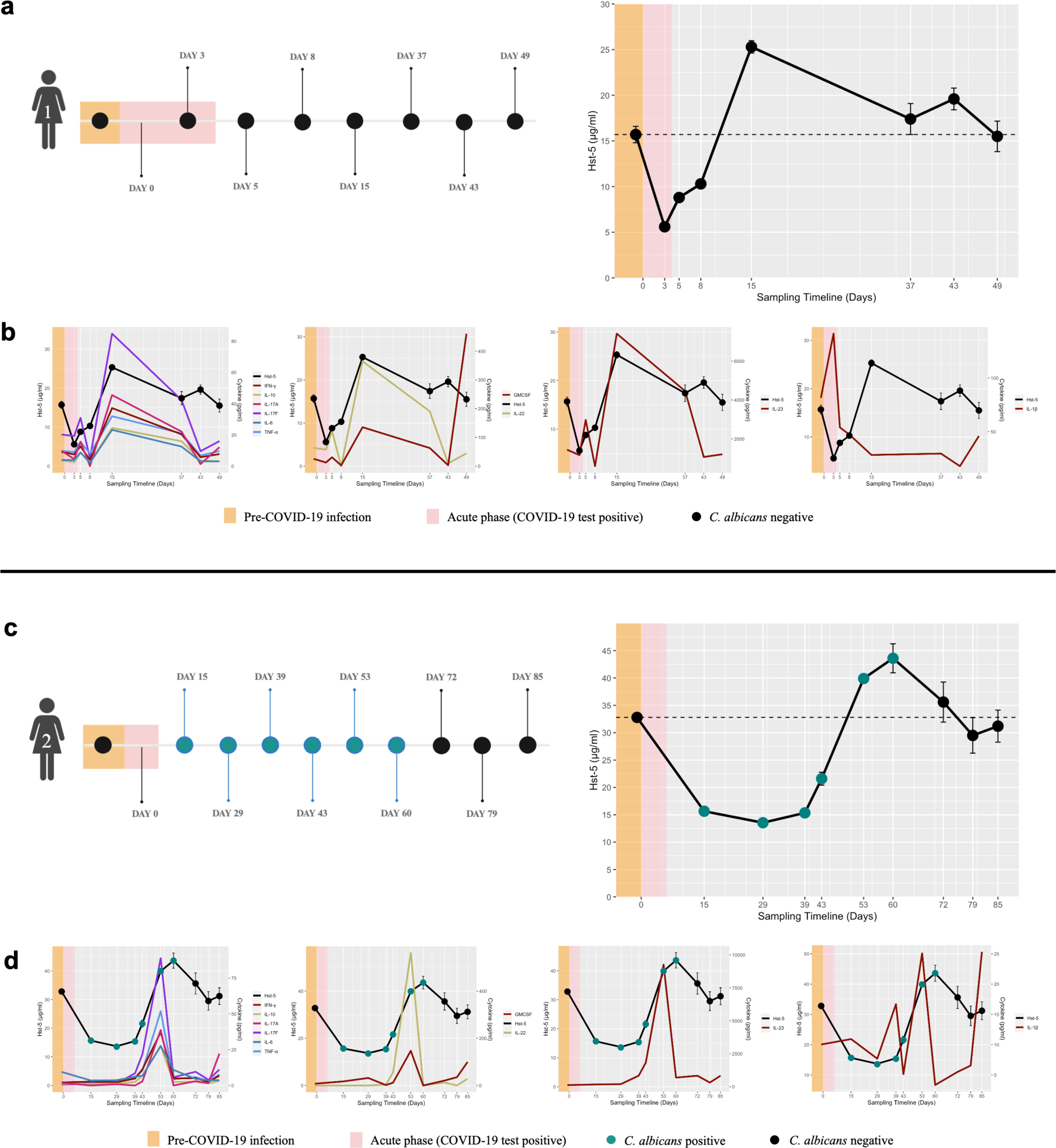
Prospective time course longitudinal analysis of saliva samples for Hst-5 from two subjects with moderate COVID-19. Changes in Hst-5 and cytokine concentrations in two longitudinally sampled subjects prior to and during the acute and post-acute phases of COVID-19 disease. Timeline of sampling during COVID-19 infection up to 49 and 85 days post-COVID-19 infection for subjects 1 and 2 **(a and c, respectively)**. Measurement of Hst-5 concentrations and fungal culturing of samples from the two subjects prior to and during the acute and post-acute phases of COVID-19 disease. Line graphs in bottom rows depict similar trends for Hst-5 and Th17 associated cytokine levels for subjects 1 and 2 **(b and d, respectively)**. Upon culturing, *C. albicans* was recovered from the initial 6 samples from subject 2 **(c, d)**.

**Figure 4.**
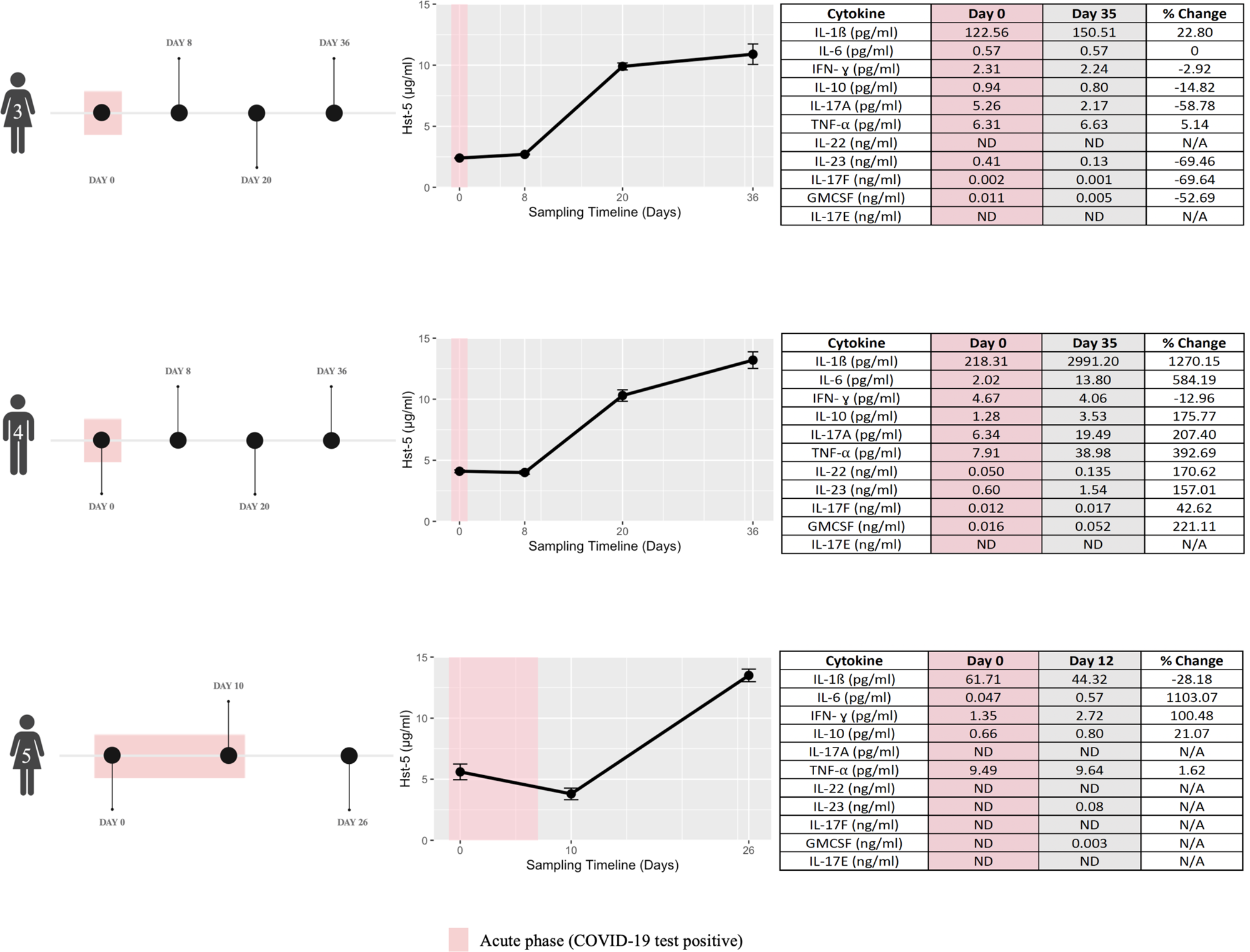
Prospective sampling and analysis of saliva from three subjects with moderate COVID-19. Timeline of sampling and measurement of salivary Hst-5 and cytokine concentrations in samples recovered during the acute phase of the disease and upon recovery. Table presents cytokine values measured in the first and last samples recovered from the three subjects and the percent change in levels between the samples.

### Activation of Th17 inflammatory pathway concomitant with changes in histatin-5 concentrations

Salivary cytokine profiling was performed on samples recovered from the 5 prospectively sampled subjects. Comparative analysis of samples from *Subject #1* demonstrated a notable increase in the Th17 associated inflammatory cytokines; compared to baseline pre-COVID-19 sample (IL-17A, IL-17F, IL-6, IL-10, TNF-⍺, IFN-*α*, IL-22, GMCSF and IL-23); concentrations gradually increased with a spike on Day 15, decreasing over subsequent days and returning to baseline level in the last sample measured (Day 39) (Fig. 3b). Although IL-1β followed the same trend, the spike in concentration was seen in the first sample recovered during the acute phase of the disease. Cytokine levels in *Subject #2* similarly demonstrated an increase in the Th17 associated inflammatory cytokines; compared to baseline pre-COVID-19 sample, concentrations of all cytokines (IL-17A, IL-17F, IL-6, IL-10, TNF-⍺, IFN-*γ*, IL-22, GMCSF, IL-23 and IL-1β) spiked on Day 38, gradually decreasing and returning to baseline level on Day 45. IL-17E was not detectable in all samples analyzed for *Subject #1* and *Subject #2* (Fig. 3b, 3d). For the remaining 3 subjects, increase in Th17-associated inflammatory cytokines was similarly observed during the post-acute phase for *Subject #3* (Fig. 4). Only the first and last samples recovered from these 3 subjects were subjected for cytokine analysis.

### *C. albicans* proliferates in saliva from COVID-19 positive subjects with low histatin-5

Saliva recovered from subjects (n=3) during acute COVID-19 infection (low histatin-5) and post-acute recovery (high histatin-5) phases were separately pooled and histatin-5 concentrations determined. Samples were subsequently tested *in vitro* for ability to control *C. albicans* proliferation (Fig. 5a). Histatin-5 concentration in the pooled saliva sample obtained during acute COVID-19 was 0.5 µg/ml (low), and in the pooled sample recovered during post-recovery concentration was 11.5 µg/ml (high) (Fig. 5b). Based on *C. albicans* growth (CFUs) following *in vitro* inoculation of saliva with 1×10^4^ cells/ml *C. albicans*, an average of 2.55×10^4^ cells/ml *C. albicans* was recovered from the low histatin-5 sample and 1.29×10^4^ cells/ml from the high histatin-5 sample (Fig. 5c).

**Figure 5.**
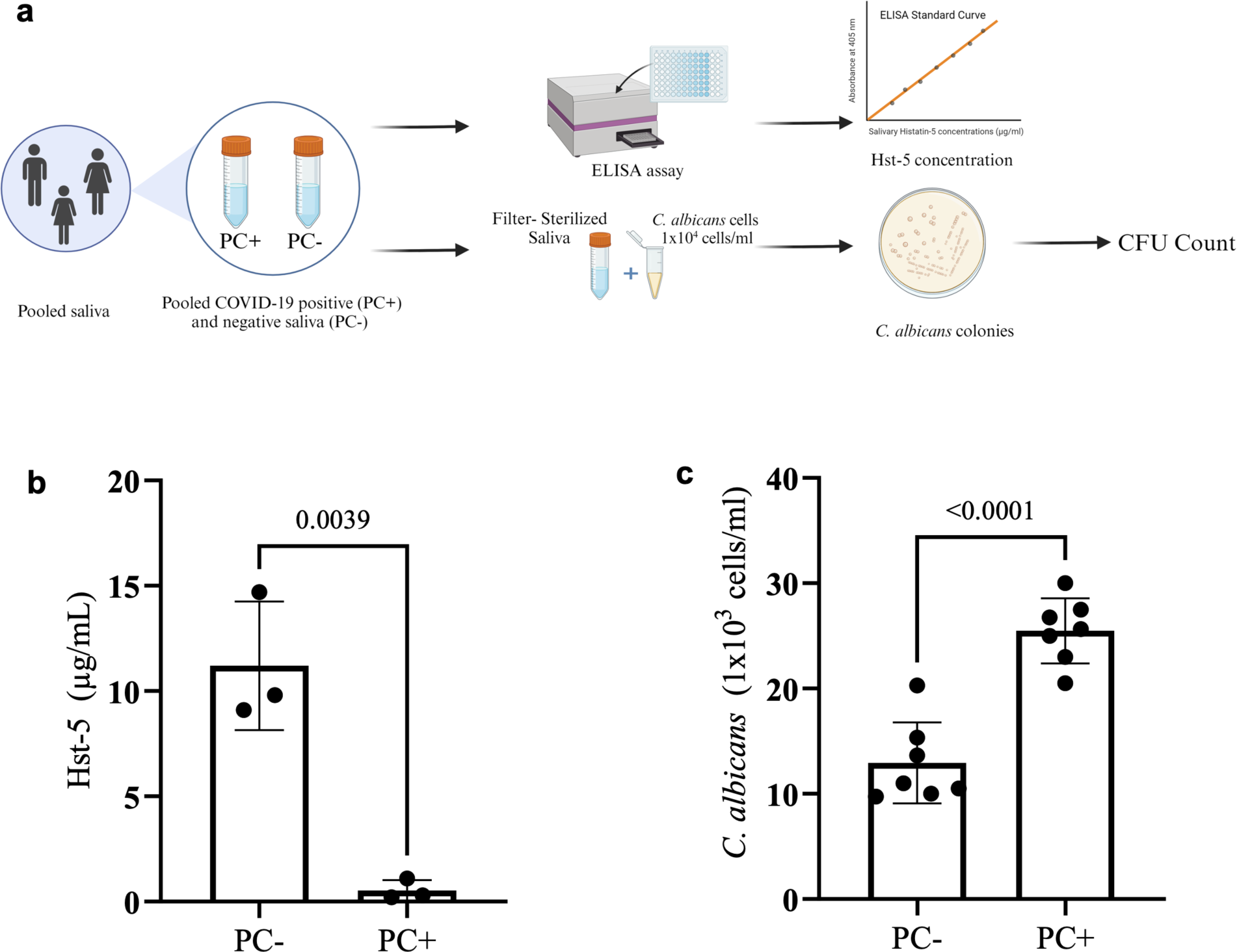
*Ex vivo* proliferation assay using pooled saliva. **(a)** Workflow for histatin-5 measurement and proliferation assay using pooled saliva samples from 3 subjects under a COVID-19 infected state (PC+) and recovered state (PC-). **(b)** Bar plots depict Hst-5 concentrations (µg/ml) from pooled saliva samples; **(c)** recovered *C. albicans* (cells/ml) following 1 h incubation in pooled saliva samples seeded with 1×10^4^ cells/ml of *C. albicans*.

## DISCUSSION

To date, little is known about the physiological mechanisms of oral manifestations in COVID-19 disease and the impact of SARS-CoV-2 infection on salivary gland function. Clinical observations are in support of SARS-CoV-2 mediated damage to the salivary glands, as COVID-19 patients frequently present with gustatory dysfunction and xerostomia, and clinical cases of inflamed salivary glands have been reported in this patient population (26).

Oral candidiasis is the most common opportunistic infection, particularly in immunocompromised individuals, although immunocompromization due to COVID-19 is ardently debated (1). However, the onset of oral candidiasis early in HIV disease indicates that immunity to *Candida* is site-specific and involves secondary local innate defenses (21, 22). In fact, innate immunity not only represents the first line of defense providing the initial host response to pathogens, but also activates the adaptive immunity (8). These systems exhibit coordinated regulation and response to establish and maintain tissue homeostasis (8). Host-produced salivary antimicrobial peptides are considered to play a vital role in innate immune defenses against microbial species, specifically histatins which are localized in the serous acinar cells and secreted into saliva (7, 12, 14).

The recent demonstration of SARS-CoV-2 replication in salivary glands and destruction of gland integrity including at sites where histatins are produced. These findings indicate that histatin production may be compromised in infected individuals. This hypothesis was validated using transcriptional analysis where the expression of both histatin genes (*HTN1* and *HTN3)* were shown to be downregulated in SARS-CoV-2 infected salivary gland tissue (Fig. 1a). Importantly, SARS-CoV-2 and histatin co-localization studies indicated an inversely proportional association where little or no histatin signal was detected in virus infected acinar cells compared to uninfected cells (Fig. 1b-f). The clinical implications of these findings were subsequently revealed using various cohorts of SARS-CoV-2 infected subjects. Overall, analysis of saliva samples demonstrated significant reduction in histatin-5 levels in infected subjects compared to healthy controls (Fig. 2). However, despite an association between infection and reduced histatin expression, the exact temporal relationship could not be perfectly gleaned. To better understand the temporal relationship between infection and histatin expression, we included a panel of prospective samples from moderately symptomatic COVID-19 subjects, which allowed us to glean some insights (Fig. 3, 4). Specifically, analysis indicated that the decrease in histatin-5 during COVID-19 is likely reversible as concentrations tended to increase during the post-acute phase of the disease. However, it is important to iterate that co-localization studies exhibited an inversely proportional association between histatin and viral counts in infected acinar cells (Fig. 1c-f), and some patients experienced long-term deficits in salivary histatin-5 levels (42 days to 1 year). Therefore, in severe COVID-19, a robust salivary infection and the ensuing adaptive immune response may irreparably damage the salivary gland parenchyma. Thus, due to pathologically lower histatin-5 secretion (and likely reduced saliva secretion overall), affected individuals may remain predisposed to both recurrent and recalcitrant to antifungal therapy oral candidiasis as part of the Long COVID syndrome (30) (https://www.covid.gov/be-informed/longcovid/about#term). It is important to note here that there are no known set “*normal*” values for salivary histatin-5 concentrations and what is considered physiological is dependent on many factors including: host factors (e.g, age, sex), time of collection, the collection method, and post-processing. Moreover, histatin-5 is expression is secreted primarily by serous acinar cells; these cells predominate in the parotid glands but are half the acinar cells in the submandibular glands, and almost entirely absent from the sublingual glands. The relative contributions of each gland to the total saliva collected is also host specific. Therefore, it is expected to see a wide range of histatin-5 concentrations among the healthy control subjects. However, based on our experience using our assay, prior clinical studies, and evaluations of anti-candidal activity, we have set arbitrary concentrations as a general guidelines for what is physiological (31).

With limited available clinical information on the hospitalized cohort population (including presence of oral lesions), we recognize that we cannot fully account for confounding factors that may influence histatin-5. However, our study focuses on the presence of SARS-CoV-2 virus rather than disease severity; importantly, on average we observed lower histatin-5 levels for infected subjects in all cohorts studied (hospitalized, outpatient and individual case). In our analysis, we did not find strong association between histatin-5 concentrations and *Candida* colonization regardless of disease status or demographics. One intriguing observation in the samples from the hospitalized cohort is the exceptionally high histatin-5 concentrations seen for 3 of the subjects with values higher than that of any of the healthy controls (Fig. 2a). Although unclear as to why, it is important to note that in the two prospectively sampled subjects with known pre-COVID-19 histatin-5 values, a spike in histatin-5 and inflammatory cytokines was noted over the course of the disease progression prior to return to baseline levels (Fig. 3). Therefore, we speculate that these three hospitalized subjects may have been sampled during these transient spikes. Interestingly, all 3 subjects with high histatin-5 levels were also positive for *C. albicans* (Fig. 2a), and it is tempting to also speculate that these high values may be part of the host immune response to *Candida* presence.

Importantly, our findings demonstrated that changes in histatin-5 concentrations were concomitant with activation of the Th17 inflammatory pathway. Antimicrobial peptides, including histatins, are part of the host Th17-type adaptive immune response (8). Interleukin-17 (IL-17) is a pro-inflammatory cytokine that regulates multiple immune events (32). In the tissues of the oral cavity, including the salivary glands, the IL-17/Th17 signaling pathway is essential for host protection against *C. albicans* infection (32, 33). Therefore, it was not surprising that cytokine profiling of saliva samples from prospectively sampled subjects displayed a trend for the Th17 associated cytokines similar to that for histatin-5 (Fig. 3). Interestingly, in the subject heavily colonized with *C. albicans*, the spike in cytokine concentrations coincided with increase in salivary histatin-5, and more significantly, with the clearance of *C. albicans* (Fig. 3c). The activation of the Th17 immune response in COVID-19 disease is expected as immunophenotyping and microscopic analysis of SARS-CoV-2 infected salivary gland tissues demonstrated chronic, focal lymphocytic sialadenitis with a T-lymphocytic infiltrates (28, 29). As IL-17 antifungal activity involves regulation of the expression of antimicrobial peptides, including histatins (8, 32, 33), the gradual increase in histatin-5 concomitant with clearance of *C. albicans* may also be mediated by activation of Th17 response. Therefore, based on these collective findings, we presume that decreased histatin-5 salivary levels due to SARS-CoV-2 infection of salivary gland serous acini permits proliferation of colonizing *C. albicans* leading to the development of oral candidiasis.

The importance of salivary histatin-5 in controlling the proliferation of *C. albicans* was shown using an *ex vivo* anti-candidal assay where saliva with low histatin-5 was able to allow *C. albicans* proliferation *in vitro* (Fig. 5). These findings provide experimental evidence establishing the importance of salivary histatin-5 in preventing *Candida* proliferation by maintaining colonizing *Candida* in the commensal state. Further, in addition to anti-candidal activity, given the established anti-inflammatory and wound healing properties of histatins (34–36), it is possible that compromised levels of histatins (and other antimicrobial proteins) may contribute to other immune- and dysbiosis-mediated mucosal conditions observed in COVID-19 patients such as burning mouth, blisters and non-specific ulcerations, and aphthous-like lesions, all of which have been associated with COVID-19 (5). With the current lack of knowledge on the implications of SARS-CoV-2 infection on oral health, the findings from this study provide mechanistic insights underscoring the oral cavities’ diverse susceptibilities to SARS-CoV-2 infection, prompting a reassessment of oral opportunistic infection risks and their potential long-term impacts on oral health.

## METHODS

### Co-localization of SARS-CoV-2 and histatin in salivary gland tissue using *in situ* hybridization and immunofluorescence, respectively

To investigate the expression of histatin during SARS-CoV-2 infection, we employed the RNAscope Multiplex Fluorescent v2 assay with antigen co-detection, adhering to the manufacturer guidelines. Studies were performed on parotid FFPE (formalin-fixed, paraffin-embedded) biopsies from uninfected donors (n=4) and from COVID-19 subjects (n=8) at autopsy with high and low copy virus number. Uninfected donors were procured from the Human Cooperative Tissue Network, and COVID-19 autopsy tissues were procured by the National Institutes of Health COVID-19 Autopsy Consortium, as described previously (28, 37). Initially, slides were dried in an air oven at 60°C for 30 min and subsequently rehydrated, and target retrieval was performed using 1X Co-Detection Target Retrieval for 20 min at 99°C. Slides were then rinsed briefly in water and washed in PBS-T (1X PBS, 0.1% Tween-20, pH 7.2), and a hydrophobic barrier was established using an ImmEdge pen. Primary antibodies against histatin-3 (Biorbyt; diluted 1:300 in Co-detection Antibody Diluent from ACD) and AQP5 (SantaCruz; diluted 1:300 in Co-detection Antibody Diluent from ACD) were applied and slides were incubated overnight at 4°C. Following washing in PBS-T, tissues and antibodies were fixed in 10% Neutral Buffered Formalin (NBF) for 30 min at room temperature, followed by another wash in PBS-T. Tissue sections were then treated with RNAscope Protease Plus for 30 min at 40°C and immediately followed by probe hybridization using either a positive control probe (320861), negative control probe (320871), or the V-nCoV2019-S probe (848561). The V-nCoV2019-S probe is antisense and specifically targets the spike protein sequence of viral RNA, providing insights into infection and viral load within the tissue. Following hybridization, signals were amplified and visualized using fluorescence Tyramine signal amplification. The primary antibodies were detected using secondary antibodies conjugated to AF488 (histatin-3), and AF568 (AQP5; aquaporin-5) and counterstaining was performed with DAPI. Images were captured using a Zeiss Axio Scan Z1 slide scanner light microscope equipped with a 40x/0.95 N.A. objective, and image files were imported into Visiopharm software version 2017.2. We first segmented the cells using the AI app from Visiopharm, once the cells were segmented, using the APP Author function we designed an algorithm that detects the dots of ISH and counts the number of dots per cell, and quantify the mean intensity of histatin-3 per cell. The data was exported and analyzed in Prism 10. Control experiments for *in situ* hybridization and immunofluorescence studies were performed prior to sample analysis (Supp. Fig. 1).

### Bulk RNAseq analysis of salivary gland tissue

Total RNA was extracted from RNAlater (Invitrogen)-preserved parotid gland at autopsy from uninfected donors (n=3) and COVID-19 subjects (n=8), using the RNeasy Mini (Qiagen) according to manufacturer protocols. Following standard bulk RNAseq using Illumina platform, normalized counts per million (CPM) were plotted for *HTN1* and *HTN3* and expression levels of both genes were compared.

### Study subjects

#### Hospitalized cohort

Adult COVID-19 patients (n=26) hospitalized at the University of Maryland Medical Center and a matched control (age, race and gender) group of healthy volunteers (n=26) were enrolled in the study. The University of Maryland Baltimore Institutional Review Board approved this study, no patient identifiers were used and informed consent was obtained from all subjects. Inclusion criteria for control subjects included: healthy adults over 18 years of age with no history of oral candidiasis, any predisposing factors or antifungal therapy. As the premise of the study is to investigate the presence of the virus in the oral cavity regardless or disease severity, the only inclusion criteria for the COVID-19 cohort was COVID-19 positivity. Of the 26 subjects, 11 were COVID-19 positive but asymptomatic and were hospitalized for other medical reasons; the remaining 15 were symptomatic with severe disease, admitted to the intensive care unit and/or receiving oxygen supplementation. The hospitalized COVID-19 population included 14 males and 12 females; 15 Caucasians and 11 African American with age range of 29-76.

#### Outpatient cohort

This cohort included 33 subjects with mild-moderate disease; a total of 68 samples were prospectively collected from these subjects at NIDCR (previously described) (28).

#### Prospective cases

In addition to the cohorts, 5 otherwise healthy outpatient subjects (4 females, one male; age range 30-58) were prospectively sampled at different timepoints during the course of their COVID-19 disease. A total of 29 samples were collected and analyzed for these subjects.

### Clinical Samples

Saliva samples were collected from participating subjects using the Salivette collection system as we previously performed (31, 38). Saliva was recovered, aliquoted and immediately stored at −80°C with protease inhibitors. In addition, oral swabs of the oral mucosal tissue were recovered for culturing to evaluate *Candida* recovery. Due to inaccessibility to some patients the result of medical status, samples for fungal culturing were only obtained from 20 of the 26 hospitalized patients. For the outpatient cohort and their control subjects, whole unstimulated saliva samples were collected and processed as above.

### Evaluation of salivary *Candida* colonization

Oral swabs from all sampled subjects were immediately cultured on fungal Yeast Peptone Dextrose (YPD) agar media (Difco Laboratories) and incubated at 35°C for 24-48 h for fungal growth evaluation. The chromogenic medium CHROMagar Candida was used for *Candida* speciation.

### Measurement of histatin-5 (Hst-5) salivary levels using ELISA

ELISA was performed as we previously described (31). A high purity Hst-5 peptide was synthesized by GenScript and Hst-5 specific rabbit polyclonal antibody was produced by Lampire Biological Laboratories. For measurement of Hst-5 levels, a standard curve was performed with each assay using Hst-5 peptide concentrations ranging from 0.5-500 μg/ml. Wells of high-binding 96-well plates were coated with 100 μl of each Hst-5 concentration or 1/100 dilution of saliva. Following overnight incubation at 4°C, wells were blocked with 0.1% dry milk in PBS for 1 h and anti-Hst-5 antibody (1/1000) (100 μl) was added for 1 h at 37°C. Following washing, HRP-labeled goat anti-rabbit secondary antibody (1/3000) (Abcam) was added, and plates incubated for 1 h at 37°C. Following washing, 100 μl of ABTS Peroxidase Substrate (KPL, Inc.) was added and plates incubated for 20 min until color develops. The reaction was stopped by the addition of 50 μl of Stop Solution (KPL, Inc.) and optical density (OD) was measured at 405nm using a microtiter plate reader. A standard curve was plotted with each run; samples were tested in triplicate on two separate occasions and the average Hst-5 concentration calculated in μg/ml. As there are no set normal Hst-5 salivary concentrations, based on our experience with clinical studies on Hst-5 salivary levels using our immunoassay, arbitrary concentrations of approximately 9-10 μg/ml was considered the cutoff whereby lower concentrations are considered in the low range (31).

### Salivary cytokines profiling

Saliva samples from outpatient subjects were analyzed at the University of Maryland Cytokine Core using the Luminex Multianalyte System. Each sample was measured in triplicate and results expressed in Pg/ml. Not enough saliva was available from other cohorts for cytokine analysis.

### *Ex vivo* salivary *C. albicans* inhibition assay

In order to assess the anti-candidal potency of saliva with respect to Hst-5 concentration, saliva samples were recovered from 3 subjects during COVID-19 and after recovery, then pooled to generate a COVID-19 positive and a COVID-19 negative sample. Hst-5 concentration in the pooled samples was pre-determined by ELISA and samples were filter-sterilized and comparatively tested for efficacy in inhibiting *C. albicans* proliferation *in vitro*. For these assays, cultures of the standard *C. albicans* SC5314 strain were grown in YPD broth (Difco Laboratories) overnight at 30°C with shaking and cells were washed and resuspended in sterile PBS (1mM). *C. albicans* cells were added to each of the saliva samples (100 μl) at final cell density of 1×10^4^ cells/ml in the wells of a 96-well microtiter plate and plates were incubated for 1 h at 37°C with shaking. Aliquots from reactions were diluted with PBS and plated on YPD agar and incubated for 24–48 h at 35°C. The number of single colonies on each plate was counted and the level of *C. albicans* proliferation was determined based on CFU counts (cells/ml).

### Statistical analysis

Statistical analysis was performed using GraphPad Prism 10.2.1. A Wilcoxon rank-sum test was used to compare histatin-5 levels between COVID-19 patients and healthy control subjects and a t-test was used to evaluate associations between histatin-5 values and patient characteristics (disease severity, *Candida* colonization, demographics). For *in vitro* assays, histatin-5 levels in pooled saliva of COVID-19 positive and negative subjects and *Candida* CFU counts were determined using the unpaired two-sample t-tests. Significance of *HTN1* and *HTN3* expression levels in parotid glands was determined using ANOVA analysis with Bonferroni correction for multiple measurement. Figures were constructed using GraphPad Prism 10.2.1 and R statistical programming software.

## ACKNOWLEDGMENT

We would like to acknowledge the contributions of LaToya Stubbs and Kendra Petrick who enrolled subjects and organized sampling of hospitalized cohort at the University of Maryland Medical Center. We would like to thank the NIH COVID-19 Autopsy Consortium (PIs-Chertow & Kleiner) for coordination and access to tissues from fatal COVID-19 cases. We give our special thanks to the members of the NIDCR Sjögren’s Disease Clinical Research Team for their coordination of patients and collection of research data and tissues. Biorender.com was used to create elements of figures.

## AUTHOR CONTRIBUTIONS

M.A.J-R., B.M.W., A.S., T.F.M., P.R. conceived and designed this research, M.A.J-R., B.M.W. provided funding; A.A.A., T.W.W., P.P., B.M.W. performed the experiments; D.E.K, D.S.C., S.M.H., B.G., S.S., S.R., collected and processed tissues and clinical data for COVID-19 autopsies; D.E.K, S.M.H., B.G., performed the autopsies; D.S.C. and D.E.K., led and oversaw the NIH COVID-19 Autopsy Consortium; M.A.J-R., A.A.A., T.W.W., P.P., B.M.W., A.S., T.F.M., analyzed data; M.A.J-R., A.A.A., T.W.W., B.M.W. wrote the paper; B.M.W. led and oversaw the execution of NIH clinical and molecular studies supporting this work; M.A.J-R. oversaw the entire study.

## ETHICAL APPROVAL

*NIH.* Autopsies are exempt from NIH single institutional review board (IRB), consent from families of fatal COVID-19 cases were obtained. Otherwise, NIH single IRB conducts ethical reviews for human research studies as required by Department of Health and Human Services regulations for the Protection of Human Subjects. All patients seen at the author’s (B.M.W.) institute (NIH/NIDCR) reported herein provided informed consent before participation in IRB-approved research protocols (NIH IRB: 20-D-0094, NCT04348240; NIH IRB: 15-D-0051, NCT02327884). Individuals on 20-D-0094 had the option to receive a $50 payment per visit ($300 total) to offset the cost of travel.

## SUPPLEMENTAL FIGURES

**Supp Figure 1. *In situ* hybridization and immunofluorescence controls. (a)** Immunofluorescence (IF) and *in situ* hybridization (ISH) controls (see also Fig. 1). Positive and negative controls for IF, antibody anti-Hst-3 in control parotid shows strong IF (green) in acini structures and no signal is observed in ducts. The isotype control shows no signal. **(b)** Probe anti-human *PPIB* shows strong signal in all the cells (white); negative control probe shows sparse, rare positive signal in the tissue.

