## Supplementary figures and images for "SARS-CoV-2 Infection of Salivary Glands Compromises Oral Antifungal Innate Immunity and Predisposes to Oral Candidiasis"

### Supplemental Figures

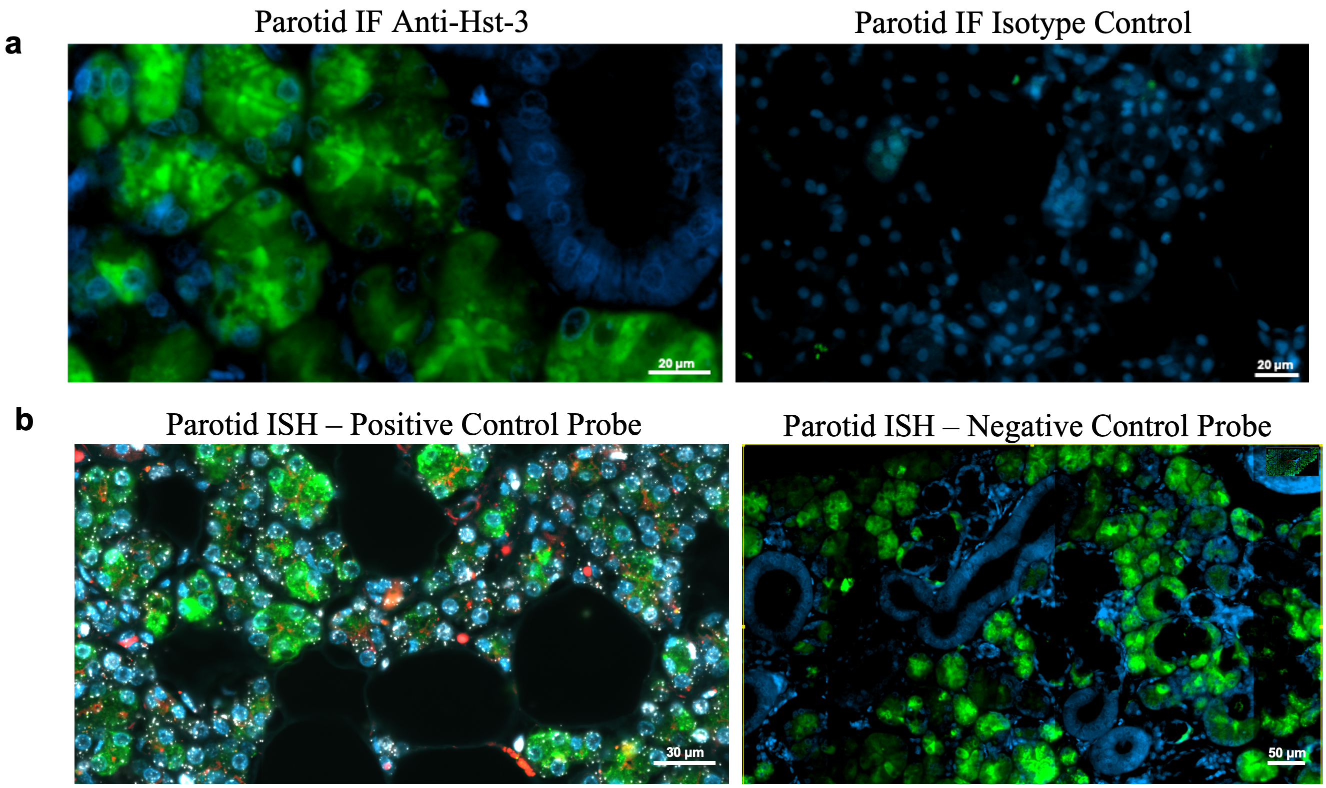
